# Structural polymorphism of *ex-vivo* ALECT2 amyloid fibrils revealed by cryo-EM

**DOI:** 10.1101/2025.07.15.664973

**Authors:** Shumaila Afrin, Binh An Nguyen, Virender Singh, Preeti Singh, Parker Bassett, Maja Pekala, Bret Evers, Christian Lopez, Yasmin Ahmed, Li Li, Raja Reddy Kallem, Andrew Lemoff, Christos Argyropoulos, Barbara Kluve-Beckerman, Lorena Saelices

## Abstract

ALECT2 amyloidosis is a rare systemic disease characterized by the pathological deposition of leukocyte cell-derived chemotaxin-2 (LECT2) as amyloid fibrils, primarily affecting the kidneys and liver. The molecular mechanisms underlying LECT2 aggregation remain poorly defined, hindering diagnostic and therapeutic development. Here, we present cryo-electron microscopy structures of *ex-vivo* ALECT2 fibrils extracted from a patient’s kidney. We identified three fibril polymorphs: a predominant single-protofilament morphology and two minor double-protofilament morphologies. The dominant single-protofilament morphology comprises the full-length 133-residue LECT2 protein and retains all three native disulfide bonds. Low-resolution reconstructions of double-protofilament morphologies suggest they adopt a similar fold to the single protofilament morphology, but form paired assemblies with different inter-filament interfaces. Mass spectrometry also reveals acetylation within the fibrils. These findings offer critical insights into the structural basis of ALECT2 amyloid formation and identify molecular features that could inform future diagnostic and therapeutic approaches.

## Introduction

ALECT2 amyloidosis is a protein disorder caused by the pathological deposition of Leucocyte Cell-Derived Chemotaxin-2 (LECT2) as amyloid fibrils in various tissues and organs^1^. The kidneys and liver are predominantly affected, while the spleen, lungs, adrenal glands, and colon are also involved to a lesser extent with varying degrees of penetrance and clinical overlap. ALECT2 amyloidosis has emerged as the third most common form of renal amyloidosis in the United States^2^. Worldwide, the disease exhibits a distinct ethnic predisposition, with higher prevalence observed in specific populations, including Hispanics of Mexican descent, Punjabis, Egyptians, Sudanese, Chinese, and Native Americans^3^. Currently, no targeted therapies are available for ALECT2 amyloidosis, and treatment remains largely supportive, with kidney transplantation considered as an option in patients experiencing advanced renal disease^2^.

The native structure of LECT2 provides insight into the factors that may contribute to its misfolding and aggregation in amyloidosis. LECT2 is produced by the liver as a 151-residue protein and released into the systemic circulation as a 133-residue protein following signal peptide cleavage. Functionally, mature LECT2 is a chemotactic factor involved in various biological processes including the immune response, where it attracts neutrophils to sites of inflammation and metabolism^4^. Structurally, the globular form of human LECT2 contains three disulfide bonds and adopts an M23 metalloendopeptidase fold. In native state, this fold includes a conserved Zn(II) coordination geometry formed by His35, Asp39, and His120, with the fourth site occupied by water instead of a coordinating residue, rendering LECT2 enzymatically inactive^4^. In disease state, the loss of zinc coordination has been suggested to destabilize the native structure of LECT2, promoting its aggregation and conversion into amyloid fibrils^5,6^. Interestingly, all patients diagnosed with ALECT2 amyloidosis are homozygous for the G-allele single-nucleotide polymorphism (at cDNA position 172; allele frequency of 0.477 overall), resulting in the presence of a valine at position 40^7-9^. *In vitro* studies using microfluidics system have shown that the LECT2-I40V variant exhibits an increased propensity for fibril formation compared to wild-type LECT2^6^.

Structural studies of amyloid LECT2 lag, with limited models largely derived from recombinant systems and computational tools. A study combining experimental and computational approaches identified two “aggregation prone regions”—residues from Val38 to Gly58 and from Val73 to Lys88, considered critical interfaces for the aggregation of recombinant LECT2^10^. A cryo-electron microscopy (cryo-EM) study of recombinant LECT2-I40V fibrils identified residues Met55 to Ile75, which forms the key intermolecular interface stabilizing the amyloid structure^11^. These observations highlight the critical need to study the structures of *ex-vivo* ALECT2 amyloid fibrils.

In the present work, we use cryo-EM to determine the structure of *ex-vivo* ALECT2 amyloid fibrils extracted from the kidney of an ALECT2 amyloidosis patient^1^. Benson and colleagues reported that this patient was homozygous for the G allele leading to I40V substitution^1^, which we confirmed by mass spectrometry. We found that ALECT2-I40V fibrils can assemble into three distinct morphologies: a single protofilament morphology (the major population) and two double protofilament morphologies (the minor populations). The single protofilament morphology incorporated all 133 residues of the LECT2 protein with the original three disulfide bonds preserved. While the density maps of the two double protofilament morphologies were of lower resolution, they appear to adopt a similar fold to the single protofilament morphology but forming paired assemblies with distinct inter-filament interfaces. Using mass spectrometry, we also detected lysine acetylation in ALECT2 amyloid fibrils. Leveraging the structural and biochemical information, we speculate on the possible mechanism of ALECT2 amyloid formation. Our study may provide critical structural information of ALECT2 amyloid fibrils, laying a foundation for developing targeted diagnostic and therapeutic approaches against ALECT2 amyloidosis.

## Results

### Histological and biochemical characterization of ALECT2 fibrils

We obtained fresh frozen ALECT2 tissue samples from the kidney of a patient homozygous for the I40V variant and performed histological staining. The patient, female, presented with nephrotic syndrome caused by renal amyloid deposits, which slowly progressed to renal failure over several years^1^. At age 61, she underwent nephrectomy for renal carcinoma, after which she progressed to end-stage renal disease requiring dialysis. The sample reported here was collected post-nephrectomy and obtained from the laboratory of Dr. Merrill D. Benson at the University of Indiana. We performed a histological analysis to confirm the presence of amyloids in kidney sections. Hematoxylin and eosin (H&E) staining highlighted tissue architecture while Congo red staining and Thioflavin-S staining of the kidney tissue sections revealed abundant amyloid deposits within the glomeruli and the interstitium of the kidney (Fig. 1a).

**Fig. 1.**
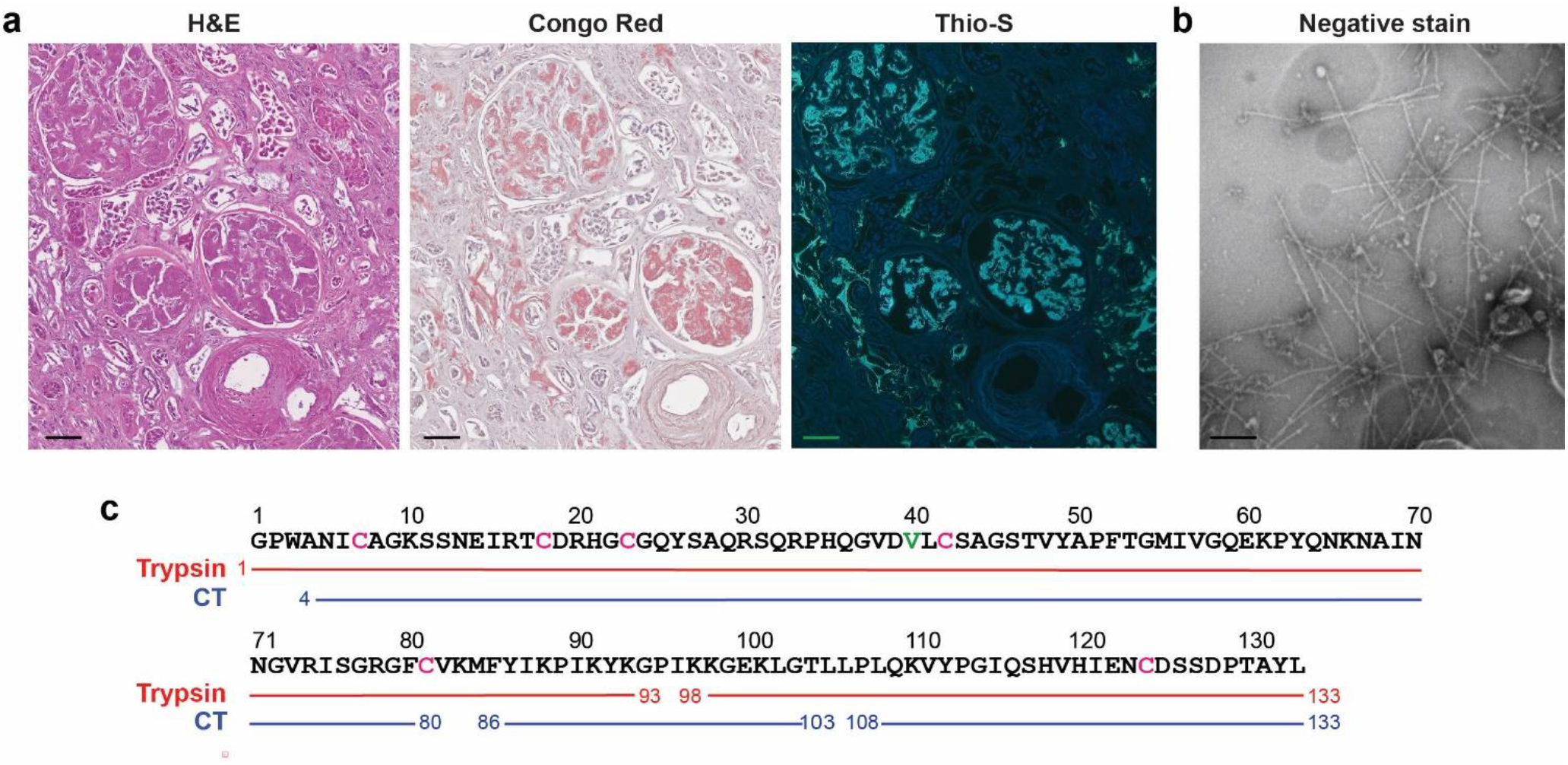
Characterization of ALECT2 amyloid deposits in kidney. **a**. Histological characterization of kidney sections using (left to right) H&E stain, Congo red, and Thioflavin-S staining. Scale bar, 50 µm. **b**. Negative stain electron microscopy image of fibrils extracted from the kidney. Scale bar, 200 nm. **c**. Proteomic analysis using tryptic (red solid line) and chymotryptic (CT, blue solid line) digestion highlights peptide coverage by mass spectrometry. Cysteines are colored pink and the Val40 is colored green.

We extracted ALECT2 fibrils from the tissue using a water-EDTA based protocol adapted from a previously described method^12^. Transmission electron microscopy with negative staining confirmed the presence of well dispersed fibrillar structures in the extractions (Fig. 1b). We verified the protein identity using mass spectrometry (Fig. 1c) and western blot analysis with a polyclonal antibody against full-length human LECT2 (Supplementary Fig. 1a). Western blot analysis showed a ∼16 KDa band that coincides with the mass of intact LECT2 protein along with additional bands at higher molecular weight, indicating the presence of multimeric species (Supplementary Fig. 1a-b). Intact mass spectrometry confirmed the presence of full-length LECT2 protein, carrying a valine at position 40 (Supplementary Fig. 1b). The intact mass spectra also revealed the presence of three disulfide bonds, with no detectable proteolytic cleavage products, consistent with the western blot analysis (Supplementary Fig. 1a-b). Interestingly, intact mass spectrometry of the fibrils revealed a secondary minor peak with an additional mass of 42 Da (Supplementary Fig. 1b and Supplementary table 1), suggestive of lysine acetylation^13^. Indeed, chymotrypsin-based mass spectrometry analysis suggests the presence of multiple lysine acetylation sites (Supplementary Table 2).

### Cryo-EM data processing, refinement, and modeling of ALECT2 amyloid fibrils

We collected cryo-EM images of ALECT2 fibrils to investigate their structural organization in the kidney (Fig. 2a). Two-dimensional (2D) classification revealed two distinct fibril morphologies: the most prevalent species consisting of one single protofilament and a minor population of fibrils consisting of double protofilaments which could further be classified into two subpopulations based on their stitching (Supplementary Fig. 2a-c). Three-dimensional (3D) classification confirmed the presence of the three morphologies (Fig. 2b). The reconstructed single protofilament morphology displayed a rise of 4.8 Å per layer and a helical crossover of 886 Å, organized with a C1 symmetry (Supplementary Fig. 3). A density map at a resolution 2.4 Å was obtained, providing detailed structural insights (Fig. 2c and Supplementary Fig. 4).

**Fig. 2.**
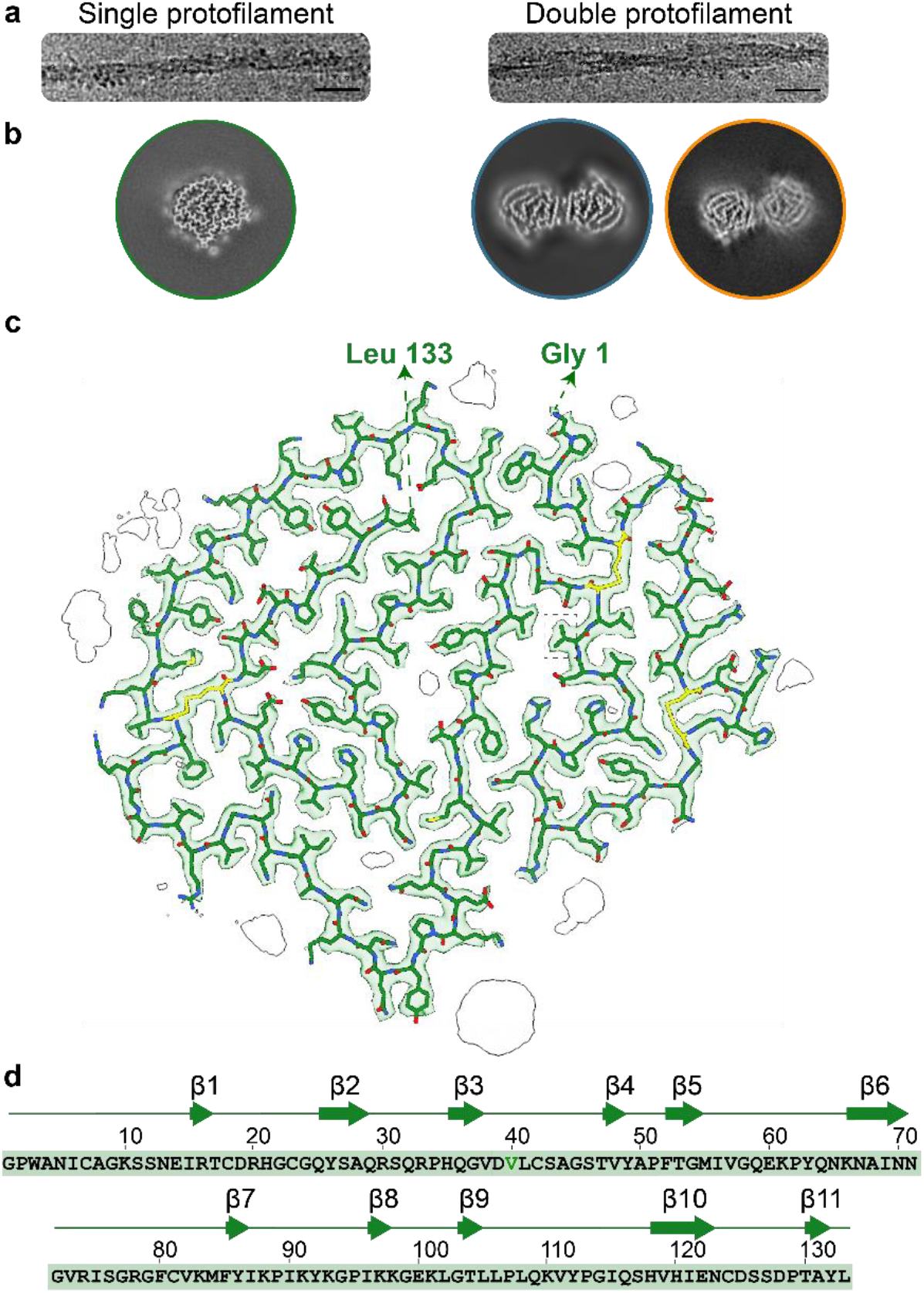
Cryo-EM analysis of ALECT2 fibrils. **a**. Representative cryo-EM micrographs showing single protofilament morphology (left) and double protofilament morphology (right). Scale bar: 20 nm. **b**. Representative 3D averages of ALECT2 fibrils. The left 3D class corresponds to the single-protofilament morphology, while the 3D classes on the right represent two distinct double-protofilament morphologies, double protofilament 1 and double protofilament 2 respectively. **c**. Single layer of ALECT2 fibril cryo-EM density map and its built-in atomic model. All 133 residues of the mature LECT2 protein are incorporated into the fibril structure, with their disulfide bonds retained (colored in yellow) and Val40 in gray dotted square. Extra densities that are predicted to be non-proteinaceous are drawn in black. **d**. Sequence of LECT2: scheme of the secondary structure elements of the single protofilament morphology. β-strands are numbered and marked with green arrows.

Leveraging the high resolution of the reconstructed map, we modeled the structure of the single protofilament morphology with precision. The accurate positioning of peptide bonds revealed that the fibrils have a left-handed twist (Supplementary Fig. 3). Additionally, the well resolved electron-density for sidechains allowed us to unambiguously validate protein identity and sequence with a valine in position 40, consistent with the mass spectrometry data (Fig. 2c-d and Fig. 1c). We modeled all 133 residues of the full-length LECT2 sequence into the amyloid structure of the single protofilament morphology, supporting the intact mass spectrometry data and western blotting (Fig. 2c and Supplementary Fig. 1). The filament fold contains eleven β-strands (Fig. 2d). The amyloid structure of the fibril retained the three original disulfide bonds bridging Cys18 with Cys23, Cys7 with Cys42, and Cys81 with Cys124 (Fig. 2c and Supplementary Fig. 5a).

### Molecular interactions stabilizing the single protofilament morphology

We observe a non-planar arrangement of layers in the single-protofilament morphology that enables inter-layer interactions within the fibril. The top view of the filament appears highly folded upon itself with Gly45 serving as an inflection point contributing to this non-planar arrangement (Fig. 3a). This bend divides the polypeptide chain into two distinct planar regions: an N-terminal segment (Gly1–Gly45) and a C-terminal segment (Ser46–Leu133) (Fig. 3a-b). The two planes are related by a 7° rotation, approximately, which enables the interaction of three consecutive layers (Fig. 3c).

**Fig. 3.**
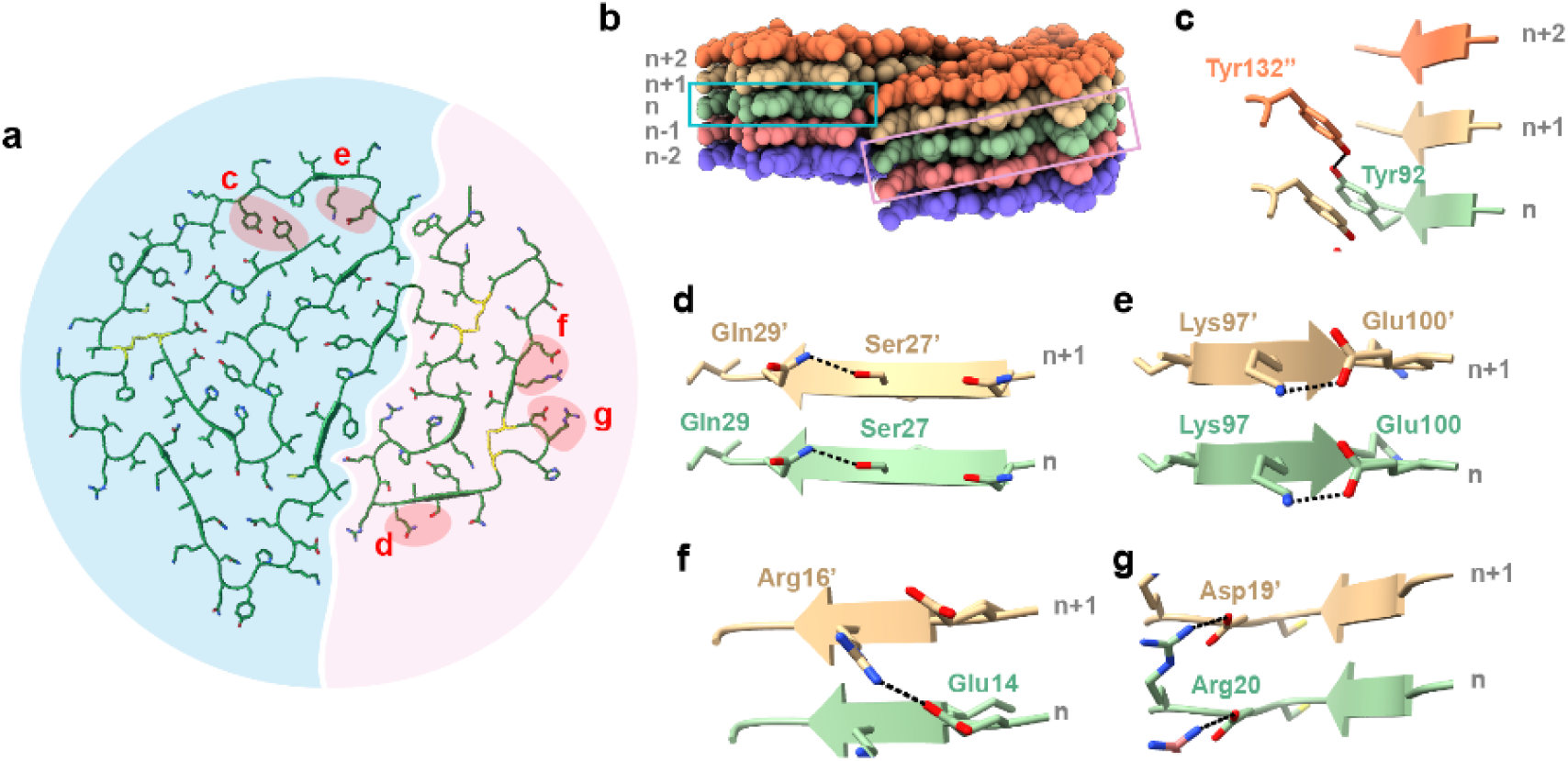
Stabilizing interactions within the ALECT2 single protofilament morphology. **a**. The top view of the fibril highlights stabilizing interactions in red; disulfide bonds are shown in yellow. The N-terminal region of the fibril, spanning residues Gly1 to Gly45, is shaded in light pink color, while the C-terminal region, encompassing residues Ser46 to Leu133, is shaded in light blue. **b**. Side view of five layers reveals an angular offset between the N- and C-terminal regions of ∼7°. **c-g**. Detailed views of examples of stabilizing interactions. **c**. Hydrogen bond between Tyr132 (*n+2*) and Tyr92 (*n*). **d**. Intra-layer hydrogen bond between Gln29 and Ser27. **e**. Salt bridge between Lys97 and Glu100 within the same layer. **f**. Salt bridge between Arg16 (*n+1*) and Glu14 (*n*). **g**. Salt bridge between Asp19 (*n+1*) and Arg20 (*n*). These interactions collectively contribute to the structural integrity of the fibril.

Additionally, we identified other molecular interactions that stabilize the fold and support fibril cohesion. These include intra-layer hydrogen bonds (Fig. 3d) and intra-layer salt bridges (Fig. 3e and Supplementary Fig. 6 a). Across the layers, the fibrils are stabilized by salt bridges such as or Glu14 of layer *n* with Arg16 of layer *n+1* or Asp19 of layer *n* with Arg20 of layer *n+1*, (Fig. 3e-f). Additionally, sidechain stacking hydrogen bonding of Asn and Gln residues form a network of polar ladders that stabilize the fibril (Supplementary Fig. 6 c-d). π-π stacking interactions by aromatic residues add further stability (Supplementary Fig. 6e)^14^. Aromatic residues Tyr92 of layer *n* also interacts with Tyr132 of layer *n+2* (Fig. 3c). The fold is further stabilized by multiple compact steric zippers formed by interdigitating residues for example residues Ile69 and Asn71 of β6 strand with residues Val119 and Ile121, of β10 strand (Supplementary Fig. 6f). Water molecules also stabilize the layers such as those interacting with the Leu106, Phe52-Thr53 and Arg16-Thr17 (Supplementary Fig. 7). In addition to the water molecules, we also observed some non-proteinaceous densities, separated from the main fibril density (Fig. 2c). These densities do not match the typical density or coordination profiles of zinc or other metal ions, and their identity remains unconfirmed. Together these interactions contribute to the overall stability of the fold and across the fibril layers. We estimated the free energy of solvation of the ALECT2 fibrils and found that their solvation energy was -67.9 kcal/mol per chain and -0.51 kcal/mol per layer. (Supplementary Fig. 8).

### Structural characterization of double protofilament morphologies

Although the number of particles was limited, we identified two distinct double-protofilament morphologies. Since the individual protofilaments in both double-protofilament morphologies adopt a fold closely resembling that of the single-protofilament morphology (Fig. 2c), we performed molecular docking to rigidly fit the single-protofilament model into the double-protofilament map (Fig. 4a). In both morphologies, the two protofilaments are inverted relative to each other. In the first morphology (C1 symmetry, 5.0 Å resolution), the inversion angle is 180, creating an interface between residues Ser11 and His21 on both protofilaments (Fig. 4a-b). In the second morphology (C1 symmetry, 5.6 Å resolution), the inversion angle is ∼35°, resulting in a smaller interface, where residues Ser11–Arg16 on one filament interact with His21–Arg16 on the other (Fig. 4a-b).

**Fig. 4.**
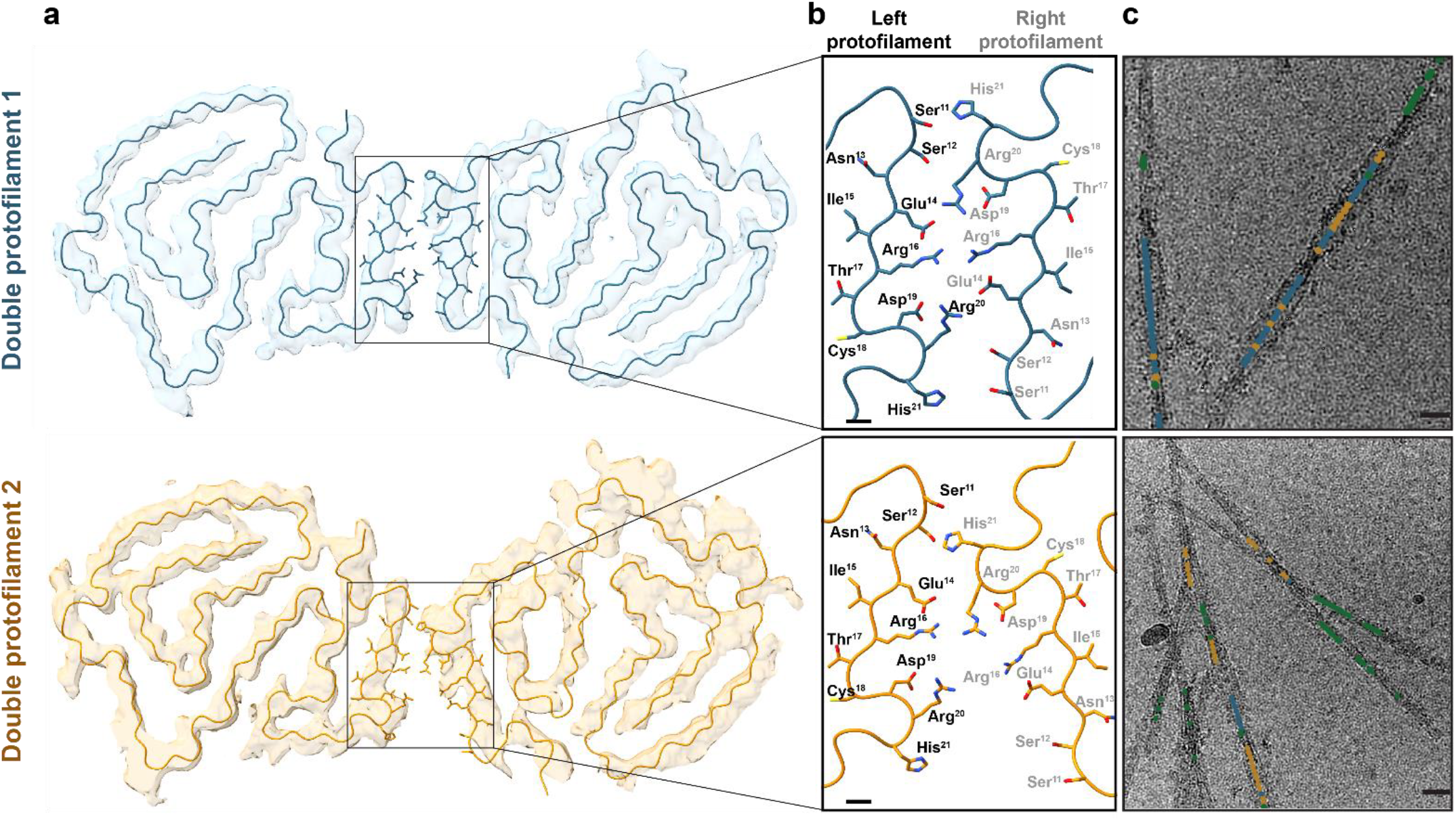
Cryo-EM reconstruction of the minor double protofilament morphologies. Cryo-EM map and docked model of the **a**. double protofilament 1 (top) double protofilament 2 (bottom) morphologies. The model for the single filament morphology was used to dock each of the two double protofilaments. **b**. Panels highlights the interface formed between the protofilaments in both morphologies. Scale bar, 3Å. **c**. Upper and lower panels are representative cryo-EM micrographs tracing the single protofilament (green), double protofilament 1 (blue) and double protofilament 2 (orange). Scale bar, 20 nm. Twenty individual micrographs were used; the representing figure is from two micrographs.

By tracing the fibril segments corresponding to these two density maps back to their micrographs, we found that both double-protofilament morphologies and the single-protofilament morphology can coexist within a single fibril (Fig. 4c). Tracing was performed using well-resolved particles that persist through the refinement process. Consequently, protofilament coverage remains incomplete and the remaining protofilaments may contain one or all three polymorphs.

### Computational prediction of Aggregation-Prone Regions (APRs)

We computationally predicted Aggregation-Prone Regions (APRs) using four prediction models (WALTZ, TANGO, Aggrescan, and AmylPred)^15-18^. We then calculated an overall APR prediction score for each residue ranging from zero to four, based on the number of prediction models that predicted the presence of an APR at that particular residue^19^. We obtained four APR predictions (Fig. 5). APR2 and APR3 were predicted by two models, and span residues Phe80 to Lys88, and Phe52 to Val57. APR1 and APR4 were predicted by only one of the four models, and span residues Val38 to Ala50, and Leu108 to Tyr112. APR1 and APR3 contain cysteine residues (Cys42 and Cys81, respectively) involved in disulfide bond formation (Fig. 5). We performed a separate analysis using the next-generation Aggrescan tool (AggresScan3D 2.0), which predicts aggregation propensity based on structural context, solvent accessibility, and amino acid composition^20^. The model identifies only two residues in the native structure as aggregation-prone, in contrast to the fibril structure, where the predicted aggregation pattern correlates with our scoring system (Fig. 5 and Supplementary Fig. 9).

**Fig. 5.**
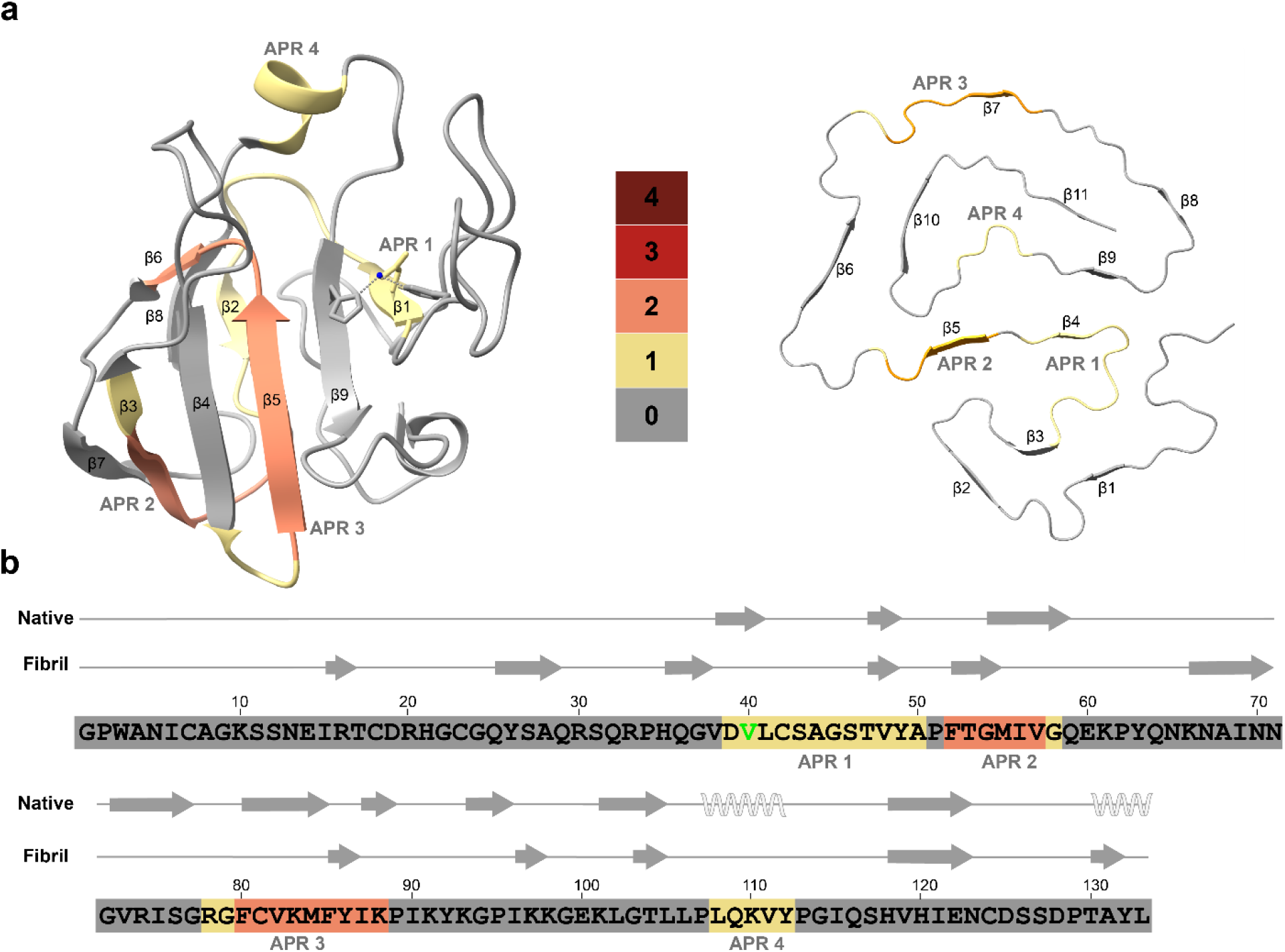
APR prediction in LECT2. **a**. Heat maps of aggregation-prone regions (APR) mapped onto the LECT2 native structure (PDB ID: 5B0H, left) and the ALECT2 single protofilament structure (PDB ID: 9NON, right) and, with scores ranging from 0 to 4 (gray to dark rust), where 0 indicates the lowest likelihood of aggregation and 4 represents the highest. **b**. ALECT2 sequence with secondary structure annotations for both the native and fibril forms, with APRs color-coded according to **a**.

## Discussion

Advances in cryo-EM resolution have provided unprecedented insights into the structural foundations of amyloid fibrils associated with amyloidosis. These structural studies have been essential in understanding the relationship between fibril architecture and disease phenotypes in many amyloid conditions^12,21,22^. This information has facilitated the development of structure-based diagnostic and therapeutic strategies^23-25^. We aim to contribute to the structural understanding of ALECT2 amyloidosis, an understudied form of systemic amyloid disease with no diagnostic and therapeutic options. In the present study, we determine and describe the structure of *ex vivo* ALECT2 fibrils, which had not been previously characterized, and discussed the implications of our findings.

Our study includes *ex-vivo* ALECT2 amyloid fibrils extracted from the kidney of a patient homozygous for I40V. We found three populations of ALECT2 fibrils, comprised of one single and two double protofilaments. We determined the cryo-EM structure of the single-protofilament ALECT2 fibrils from the kidney, and found a unique architecture, with the full-length LECT2 protein forming the amyloid core, retaining native disulfide bonds, and incorporating post-translational acetylation modifications (Fig. 2 and Supplementary Fig. 1). Additionally, we obtained low-resolution density reconstructions of two double-protofilament morphologies from the same patient and tissue (Fig. 2b and Fig. 4). Each protofilament within these double-protofilament morphologies closely resembles the single protofilament morphology. The two morphologies differ in their inter-filament interface (Fig. 4).

The ALECT2 single-protofilament morphology is likely stabilized by multiple factors. Unlike most other amyloidogenic proteins to date, where only specific regions of the protein form the amyloid core, the entirety of the LECT2 sequence is incorporated into the fibril core (Amyloid Atlas 2025)^26^. This observation is consistent with previous case studies that reported the full-length sequence of LECT2 incorporated into ALECT2 amyloid fibrils, using mass spectrometry^1^. It remains unclear whether the incorporation of full-length LECT2 into the amyloid core confers resistance to proteolysis, or whether resistance to proteolysis facilitates the formation of a full-length amyloid core. Nevertheless, the end result may be an enhanced fibril stability, although this hypothesis remains to be validated through *in vitro* experiments. Computational solvation energy calculations, previously employed as a proxy for structural stability^14^, suggest that the presence of full-length LECT2 in the fibril core may indeed contribute to the overall high stability. While ALECT2 fibrils have a per-residue solvation energy of -0.51 kcal/mol, the incorporation of all 133 amino-acid residues results in a total estimated solvation energy of -67.9 kcal/mol per molecule (Supplementary Fig. 8). This value falls at the lower end of the current range reported for amyloid fibrils, suggesting high fibril stability (Amyloid Atlas, 2025)^26^. Additional factors that could contribute to this stability may arise from a combination of structural features, including intralayer disulfide bonds, an extensive hydrogen-bonding network, and salt bridges within and across fibril layers (Fig. 3 and Supplementary Fig. 6a-e). Additionally, the ALECT2 fibril core is enriched with steric zippers, which provide further structural reinforcement by tightly interlocking adjacent fibril strands (Fig. 3 and Supplementary Fig. 6f)^14^.

Unlike other amyloid proteins^19,27^, LECT2 is not predicted to be highly amyloidogenic. Only two of four prediction tools (AGGRESCAN and AmylPred) identify aggregation-prone regions (APRs) within the protein sequence (Fig. 5). Moreover, the structure-based model AggresScan3D 2.0 predicts aggregation propensity at only two residues in the native LECT2 structure (Supplementary Fig. 9). These findings raise questions about how APRs contribute to aggregation or influence fibril stability. It is possible that zinc loss, local pH disturbances, shearing forces, and/or post-translational modifications destabilize the native fold of LECT2, exposing these APRs, and enabling aggregation. Drawing on insights from the cryo-EM structure, mass spectrometry, and amyloid propensity analyses, we speculate on several potential contributors and requirements for amyloid fibril formation in ALECT2 amyloidosis, as explored further below.

Post-translational modifications have been proposed to influence protein aggregation in amyloid conditions. Currently, no published reports exist on post-translational modifications of native LECT2 protein or ALECT2 amyloid fibrils. However, our intact mass data, combined with post-translation modification analysis using chymotrypsin-based mass spectrometry, identified lysine acetylation (Nεacetylation) in ALECT2 fibrils (Supplementary Table 1). The functional implications of this acetylation remain unclear and are likely context dependent. In other amyloid systems, the effects of lysine acetylation on aggregation vary significantly. For instance, acetylation at Lys16 in Aβ42 reduces aggregation propensity^28^, while in tau, acetylation of lysines adjacent to amyloidogenic regions have more variable effects on tau aggregation, some inhibiting and some promoting^29,30^. Acetylation of lysines could promote ALECT2 fibril formation by destabilizing the native structure of LECT2 or by stabilizing the structure of fibrils. Although recombinant studies show that acetylation may not be required for LECT2 aggregation^11^, acetylation at certain positions of the native form of LECT2 could interfere with the native hydrogen bonding between the side-chain amide from the lysine and the backbone carbonyl oxygen from neighboring prolines, thereby destabilizing the native structure (Supplementary Fig. 10). In ALECT2 fibrils, acetylated lysines, which are solvent exposed (Supplementary Table 2 and Supplementary Fig. 8), could neutralize side chain charges, thereby influencing fibril structure stability, their interactions within the fibrils or with other proteins, and/or prevent clearance^29^. However, due to the limited experimental data available, it remains challenging to determine the timing of acetylation or its specific role in ALECT2 amyloidosis. Further exploration is necessary to understand how this modification influences protein-protein interactions and whether it plays a functional role in the pathogenesis of ALECT2 amyloidosis.

A critical early event may involve the loss of zinc binding, either due to reduced zinc levels or pH changes, destabilizing the native LECT2 structure. Our *ex vivo* ALECT2 amyloid structure revealed a complete loss of the zinc-binding pocket formed by Asp39 and His120 (Supplementary Fig. 8). This observation aligns with prior studies identifying zinc dissociation as a key trigger for ALECT2 amyloidogenesis^5,6^. The resulting destabilization could initiate a cascade of structural rearrangements, particularly affecting the protein’s secondary structure elements, as suggested by comparing the β-strands in the native and fibril structures (Fig. 5b). The I40V mutation is believed to increase the conformational heterogeneity of the zinc free form of LECT2, rather than facilitating zinc loss^5^. Despite the high allele frequency of I40V in some populations (approximately 50% in Hispanics, for instance), the disease prevalence is considerably lower, suggesting that additional factors may influence amyloidogenesis or that the disease is extremely underdiagnosed^6^.

Fibril formation in ALECT2 amyloidosis requires partial or complete unfolding of the native protein. Intact mass spectrometry (Supplementary Fig. 1) and the cryo-EM density map (Fig. 2) confirm that the disulfide bonds remain intact, as observed in ALys and AL amyloid fibrils^19,31^. However, detailed analysis of the cryo-EM structure reveals that two disulfide bonds, Cys7–Cys42 and Cys81–Cys124, undergo a reorientation (Supplementary Fig. 5). This structural rearrangement likely requires at least partial unfolding of the native protein, exposing buried APRs and promoting fibril formation. Since the disulfide bonds are preserved, misfolding may take place in an oxidative environment that prevents their reduction, such as the extracellular space or the endocytic/secretory pathway^32,33^. Unlike ATTR amyloidosis, where circulating transthyretin aggregates have been detected in plasma^23^, this observation suggests that LECT2 aggregation may occur locally within affected tissues. Local factors such as zinc loss, acidic pH, oxidative stress, or extracellular matrix interactions may facilitate native-state destabilization and drive the transition toward amyloid formation.

Comparison between the structures of *ex-vivo* ALECT2 fibrils and recombinant LECT2 fibrils suggests a potential influence of the environment in which fibrils are formed. Although the recombinant fibrils were generated from the full-length protein, their structure substantially differs from the *ex-vivo* ALECT2 fibrils reported here (Supplementary Fig. 11)^11^. The primary distinction lies in the fibril core: recombinant fibrils contain a short segment spanning residues 55–75^11^, rather than the full-length protein observed in *ex-vivo* fibrils. Given that the sequences of the precursor proteins are identical, this structural disparity likely arises from differences in environmental or biochemical conditions during fibril formation. Previous studies suggest that shear forces, such as those present in kidney vessels, may trigger LECT2 aggregation and promote the integration of the full-length protein into the fibril core^6^. Nonetheless, the structural similarity observed between residues 62–71 of the recombinant and *ex-vivo* fibrils (Supplementary Fig. 11) indicates that local sequence characteristics may also influence fibril formation and structural assembly. Additional studies are required to elucidate the precise mechanisms underlying LECT2 aggregation.

## Conclusions

In summary, this study highlights the structural polymorphism of *ex-vivo* ALECT2 amyloid fibrils and presents a high-resolution structure of ALECT2 fibrils, revealing one single protofilament with all 133 amino acid residues incorporated into the amyloid core. Additionally, we obtained low-resolution density maps of double protofilament morphologies, which have similar fold to each other and to the single protofilament but differ in their inter-filament interfaces. We speculate on the mechanism of ALECT2 aggregation involving zinc ion loss and partial protein unfolding. Our findings raise important questions about the role of post-translational modifications and other contributing factors in ALECT2 pathogenesis. Identifying these factors will be crucial for fully understanding disease progression and warrants further investigation. Targeting the *ex-vivo* ALECT2 structure may offer novel targets for detection and treatment of ALECT2 amyloidosis.

## Methods

### Patients and tissue material

Frozen kidney tissue specimen from an ALECT2 amyloidosis patient^1^ were obtained from the laboratory of the late Dr. Merrill D. Benson at the University of Indiana. The Office of the Human Research Protection Program granted exemption from the Internal Review Board review because the specimens were anonymized.

### Histology

Fresh-frozen tissue was thawed at 4°C overnight and then fixed in 20X volume excess of 10% neutral buffered formalin for 48-hours at room temperature with agitation. Then, samples were transferred to 70% ethanol and paraffin processed according to established protocols^34^. Sections from prepared paraffin blocks were prepared for routine hematoxylin and eosin staining (H&E), Congo Red and Thioflavin-S staining. Regressive H&E was performed on a Sakura DRS-601 x,y,z robot utilizing Leica-Surgipath Selectech reagents following the Sheehan’s textbook methodology. Congo Red slides counterstained with hematoxylin were evaluated for pathologic amyloid aggregation under bright-field. Color was imparted to amyloid by utilizing 0.1% Congo Red in alcoholic-saline following sensitization of the slides with alkaline-alcoholic-saline (10% NaCl/0.5% NaOH/80% EtOH). Slides stained with Thioflavin-S were examined for amyloid deposits indicative of plaque-like and vascular amyloid. These deposits emit bright fluorescence upon excitation with ultraviolet light (400–440 nm excitation, emission captured using a 470 nm+ long-pass filter). The staining procedure followed an adapted protocol originally detailed by Guntern et al^35^. Briefly, paraffin-embedded tissue sections were first hydrated and subjected to a series of chemical pretreatments: initial oxidation with potassium permanganate (0.25% w/v), followed by bleaching steps with potassium metabisulfite (1%) and oxalic acid (1%), peroxidation using hydrogen peroxide (1%) in sodium hydroxide solution (2%), and finally acidification with acetic acid solution (0.25%). Each chemical step was interspersed with thorough water rinses. Subsequently, slides were equilibrated in 50% ethanol and stained for seven minutes in a solution of Thioflavin-S (0.004%) in ethanol. After staining, excess dye was removed by washing in ethanol solutions, followed by dehydration and clearing. Finally, slides were cover-slipped using Cytoseal 60, a non-fluorescent permanent mounting medium (Epredia, Kalamazoo, MI).

### Extraction of ALECT2 amyloid fibrils

ALECT2 amyloid fibrils were isolated from fresh-frozen human tissue following a previously established protocol with modifications^12^. Approximately 300 mg of frozen tissue was first thawed at room temperature and finely minced using a scalpel. The tissue fragments were then suspended in 0.5 mL of Tris-calcium buffer (20 mM Tris, 138 mM NaCl, 2 mM CaCl_2_, 0.1% NaN_3_, pH 8.0) and centrifuged at 7000 × g for 5 minutes at 4 °C. The resulting pellet underwent four additional washes with the same Tris-calcium buffer. Following these washes, the pellet was incubated overnight at 37 °C with agitation at 700 rpm in 0.5 mL collagenase solution (5 mg mL^−1^ collagenase dissolved in Tris-calcium buffer) containing cOmplete EDTA-free protease inhibitor cocktail (Roche). After incubation, the mixture was centrifuged at 7000 × g for 30 minutes at 4 °C. The pellet obtained was then resuspended in 0.5 mL of Tris–EDTA buffer (20 mM Tris, 140 mM NaCl, 10 mM EDTA, 0.1% NaN_3_, pH 8.0), followed by centrifugation at 7000 × g for 5 minutes at 4°C. This washing step with Tris–EDTA buffer was repeated nine more times. After completing these washes, fibrils were extracted by resuspending the pellet in 150 μL ice-cold water supplemented with 5 mM EDTA and centrifuging at 3100 × g for 5 minutes at 4°C. The fibril extraction procedure was repeated twice more.

### Negative-stained transmission electron microscopy

The presence of ALECT2 amyloid fibril was confirmed by transmission electron microscopy as described^12^. Briefly, a 4 μL fibril sample was spotted onto a freshly glow-discharged carbon film 300 mesh copper grid (Electron Microscopy Sciences), incubated for 1 min, and gently blotted onto a filter paper to remove the solution. 4 μL uranyl acetate was applied onto the grid and immediately removed. The grid was negatively stained with 4 µL of 2% uranyl acetate for 1 min and gently blotted to remove the solution. A FEI Tecnai G2 Spirit Biotwin electron microscope (Thermo Fisher Scientific) at an accelerating voltage of 120 kV was used to examine the specimens.

### Western blot of extracted ALECT2 fibrils

Western blot was performed on the extracted fibrils. Briefly, 0.5 µg of fibrils were dissolved in tricine SDS sample buffer and boiled for 2 minutes at 85 °C. The samples were then loaded and run onto a Novex™ 16% tris-tricine gel system using a tricine SDS running buffer. After electrophoresis, the proteins were transferred to a 0.2 µm nitrocellulose membrane. The membrane was probed with a polyclonal goat anti-human antibody targeting full length human LECT2 protein (Cat No. AF722; Lot: CZB0421051; R&D Systems™, 1:1000). A horseradish peroxidase-conjugated rabbit anti-goat IgG (Cat No. SA00001-4; Proteintech Group Inc, 1:1000) was used as the secondary antibody. The LECT2 protein was visualized using Promega Chemiluminescent Substrate, following the manufacturer’s instructions.

### Mass Spectrometry (MS) sample preparation, data acquisition and analysis

<u>For tryptic and chymotryptic MS analysis</u>, 0.5 µg of extracted fibrils were dissolved in a tricine SDS sample buffer, boiled for 2 minutes at 85°C, and run on a Novex™ 16% tris-tricine gel system using a Tricine SDS running buffer. Gel was stained with Coomassie dye, destained and ALECT2 smear was cut from the gel. Sample was sent for MS analysis. Samples were digested overnight with trypsin (Pierce) or chymotrypsin following reduction and alkylation with DTT and iodoacetamide (Sigma–Aldrich). The samples then underwent solid-phase extraction cleanup with an Oasis HLB plate (Waters) and the resulting samples were injected onto a Q Exactive HF mass spectrometer coupled to an Ultimate 3000 RSLC-Nano liquid chromatography system. Samples were injected onto a 75 um i.d., 15-cm long EasySpray column (Thermo) and eluted with a gradient from 0-28% buffer B over 90 min. Buffer A contained 2% (v/v) ACN and 0.1% formic acid in water, and buffer B contained 80% (v/v) ACN, 10% (v/v) trifluoroethanol, and 0.1% formic acid in water. The mass spectrometer operated in positive ion mode with a source voltage of 2.5 kV and an ion transfer tube temperature of 300°C. MS scans were acquired at 120,000 resolution in the Orbitrap and up to 20 MS/MS spectra were obtained in the ion trap for each full spectrum acquired using higher-energy collisional dissociation (HCD) for ions with charges 2-8. Dynamic exclusion was set for 20 s after an ion was selected for fragmentation.

Raw MS data files were analyzed using Proteome Discoverer v3.0 SP1 (Thermo), with peptide identification performed using a semitryptic or semichymotryptic search with Sequest HT against the human reviewed protein database from UniProt. Fragment and precursor tolerances of 10 ppm and 0.02 Da were specified, and three missed cleavages were allowed. Carbamidomethylation of Cys was set as a fixed modification, with oxidation of Met set as a variable modification. The false-discovery rate (FDR) cutoff was 1% for all peptides.

For <u>intact protein mass analysis</u>, 20 µL (5 µg) of extracted ALECT2 fibrils was spun at 21000 x g for 1h at 4 °C. The 10 µL of supernatant was removed and 5 µL of 8M GuHCl was added to dissociate the fibrils. Mixture was incubated at 37 C for 1 at with continuous shaking at 1000 rpm (Thermomizer). Then, an additional 5 µL 8M GuHCl was added and again incubated at 37°C for 1 at with continuous shaking at 1000 rpm. Samples with or without reducing agent (DTT) were desalted and analyzed by LC/MS, using a Sciex X500B QTOF mass spectrometer coupled to an Agilent 1290 Infinity II HPLC. Samples were injected onto a POROS R1 reverse-phase column (2.1 × 30 mm, 20 µm particle size, 4000 Å pore size) and desalted. The mobile phase flow rate was 300 µL/min and the gradient was as follows: 0-3 min: 0% B, 3-4 min: 0-15% B, 4-16 min: 15-55% B, 16-16.1 min: 55-80% B, 16.1-18 min: 80% B. The column was then re-equilibrated at initial conditions prior to the subsequent injection. Buffer A contained 0.1% formic acid in water and buffer B contained 0.1% formic acid in acetonitrile.

The mass spectrometer was controlled by Sciex OS v.3.0 using the following settings: Ion source gas 1 30 psi, ion source gas 2 30 psi, curtain gas 35, CAD gas 7, temperature 300 °C, spray voltage 5500 V, declustering potential 135 V, collision energy 10 V. Data was acquired from 400-2000 Da with a 0.5 s accumulation time and 4 time bins summed. The acquired mass spectra for the proteins of interest were deconvoluted using Bio Tool Kit within Sciex OS in order to obtain the molecular weights. Peaks were deconvoluted over the entire mass range of the mass spectra, with an output mass range of 7000-9000 Da, using low input spectrum isotope resolution.

For<u>post-translation modification analysis</u>, 0.5 µg of extracted fibrils were dissolved in a tricine SDS sample buffer, boiled for 2 minutes at 85 °C, and run on a Novex™ 16% tris-tricine gel system using a Tricine SDS running buffer. Gel was stained with Coomassie dye, destained and ALECT2 smear was cut from the gel. Sample was sent for MS analysis. Samples were digested overnight at room temperature with chymotrypsin (Worthington) following reduction and alkylation with DTT and iodoacetamide (Sigma–Aldrich). Following solid-phase extraction cleanup with an Oasis HLB µelution plate (Waters), the resulting peptides were reconstituted in 10 uL of 2% (v/v) acetonitrile (ACN) and 0.1% trifluoroacetic acid in water. 5 uL of each sample were injected onto an Orbitrap Fusion Lumos mass spectrometer (Thermo Electron) coupled to an Ultimate 3000 RSLC-Nano liquid chromatography system (Dionex). Samples were injected onto a 75 μm i.d., 75-cm long EasySpray column (Thermo), and eluted with a gradient from 0-28% buffer B over 90 min. Buffer A contained 2% (v/v) ACN and 0.1% formic acid in water, and buffer B contained 80% (v/v) ACN, 10% (v/v) trifluoroethanol, and 0.1% formic acid in water. The mass spectrometer operated in positive ion mode with a source voltage of 2.5 kV and an ion transfer tube temperature of 300 °C. MS scans were acquired at 120,000 resolution in the Orbitrap and up to 10 MS/MS spectra were obtained in the Orbitrap for each full spectrum acquired using higher-energy collisional dissociation (HCD) for ions with charges 2-7. Dynamic exclusion was set for 25 s after an ion was selected for fragmentation.

Raw MS data files were analyzed using Proteome Discoverer v3.0 SP1 (Thermo), with peptide identification performed using Sequest HT searching against the human reviewed protein database from UniProt. Fragment and precursor tolerances of 10 ppm and 0.02 Da were specified, and up to six missed cleavages were allowed with cleavage allowed at Phe, Leu, Met, Trp, and Tyr. Carbamidomethylation of Cys was set as a fixed modification, with phosphorylation of Ser, Thr, Tyr, acetylation of Lys, methylation of Lys and Arg, dimethylation of Lys and Arg, and trimethylation of Lys were set as variable modifications. The false-discovery rate (FDR) cutoff was 1% for all peptides.

The mass spectrometry proteomics data has been deposited to MassIVE (a member of ProteomeXchange)^36^ and can be accessed at MSV000098220

### Cryo-EM sample preparation, data collection, and processing

Freshly extracted fibril samples (3 µL) were applied to glow-discharged R 1.2/1.3, 300 mesh, Cu grids (Quantifoil), blotted with filter paper to remove excess sample, and plunge-frozen in liquid ethane using a Vitrobot Mark IV (FEI/Thermo Fisher Scientific). Cryo-EM samples were screened on Talos Arctica at the Cryo-Electron Microscopy Facility (CEMF) at The University of Texas Southwestern Medical Center (UTSW), and the final datasets were collected on a 300 keV Titan Krios microscope (FEI/Thermo Fisher Scientific) with ColdFEG/Falcon4i/Selectris operated at slit width of 10 eV at the Stanford-SLAC Cryo-EM Center (S^2^C^2^). Pixel size, frame rate, dose rate, final dose, and number of micrographs per sample are detailed in (Supplementary Table 3). The raw movie frames were gain-corrected, aligned, motion-corrected and dose-weighted using RELION’s own implemented motion correction program^37^. Contrast transfer function (CTF) estimation was performed using CTFFIND 4.1^38^. All steps of helical reconstruction, three-dimensional (3D) refinement, and post-process were carried out using RELION 4.0^39,40^. The filaments were picked automatically using Topaz in RELION 4.0^41,42^. Particles were extracted using a box size of 1024 and 300 pixels with an inter-box distance of 3 asymmetrical units at helical rise of 4.8 Å. 2D classification of 1024-pixel particles was only used to estimate the fibril crossover distance. 2D classifications of 300-pixel particles were used to remove suboptimal segments. An initial 3D reference model was generated from a subset of 2D class averages using relion_helix_inimodel2d program, as previously described^43^. Fibril helix was assumed left-handed for 3D reconstruction and 3D auto refinements were performed. Subsequent 3D auto refinements with optimization of helical twist and rise were carried out once the estimated resolution of the map reached beyond 4.75 Å. 3D classifications without particle alignment were used to further remove suboptimal segments, and to separate different conformation as in the case of the double filaments. Particles potentially leading to the best reconstructed map were further chosen and run through additional 3D auto-refinements. CTF refinements, 3D auto refinements, and post-processing were repeated to obtain higher resolution. The final overall resolution was estimated from Fourier shell correlations at

0.143 threshold between two independently refined half-maps (Supplementary Fig. 4).

### Atomic model building and refinement

We used the automated machine-learning ModelAngelo approach with minor modifications to obtain an initial atomic model^44^. First, COOT v0.9.8.1 was used to build the peptide backbone of a single fibril layer featuring all alanine residues^45^. We then created a new density map of one fibril layer in ChimeraX v1.8 using the command ‘vol zone #1 near #2 range 3 new true’, where #1 refers to the post-processed fibril density map and #2 is the peptide backbone model^46^. This new map was then entered into ModelAngelo, both with and without the primary LECT2 sequence, to obtain the initial atomic models of ALECT2 fibrils. Using COOT, we made residue modifications and real-space refinements to finalize the model. Further refinement was carried out using ‘phenix.real_space_refine’ from PHENIX 1.20^47^. ChimeraX v1.8 was used for molecular graphics and structural analysis^48^. Model statistics are summarized in Supplementary Table 3.

### Particle tracing

Particles used in helical reconstruction were selected from the output of the 2DClass jobs as class averages. Class averages from RELION corresponding to the desired morphology were selected using subset selection, which resulted in a particles.star file for each particular subset or morphology. We used class averages with a resolution better than 5 Å. This cutoff balances a suitable number of particles per micrograph while maintaining a high confidence that the class averages match the intended morphology. Using the CoordinateX and CoordinateY columns within the particles.star file, each individual particle was graphed as a scatterplot and then overlaid onto the corresponding micrograph listed in the MicrographName column. Please note that only clearly resolved segments of fibrils were utilized in the reconstruction process, while segments of insufficient quality were excluded. Consequently, the selected segments, even when derived from a single fibril, might appear discontinuous or scattered, as depicted in Fig. 4c.

### Stabilization energy calculation

The stabilization energy per residue was calculated by the sum of the products of the area buried for each atom and the corresponding atomic solvation parameters (Supplementary Fig. 8). The overall energy was calculated by the sum of energies of all residues, and different colors were assigned to each residue, instead of each atom, in the solvation energy map^14^.

### APR analysis

Four prediction models; TANGO^16^, WALTZ^17^, Aggrescan^15^, and AmylPred^18,49^ were used to predict APRs. The LECT2 sequence was input to a file. Residues identified as APRs by a model (“hits”) received a score of 1, while non-APR residues received a score of 0. This process was repeated across all four models, and the per-residue score was calculated as the sum of hits from the models. Criteria for calling hits were based on the requirements set by the creators of each model. Briefly, they are as follows: For TANGO, hits were counted for any stretch of 5 residues or more with a 5.0 or higher aggregation score, according to the TANGO usage guidelines. For WALTZ, hits were counted for any residue with a score greater than 0. For Aggrescan, hits were counted as any stretch of 5 or more residues with an aggregation score of 0.02 or higher. For AmylPred, hits were counted as any residues with a consensus score of 3 or more. The cumulative hits for each residue were tabulated and recorded in a PDB file as B-factors, enabling visualization in a heatmap model. The heatmap was generated using ChimeraX^46^ (v1.7.1.4). For Aggrescan 3D 2.0, PDBs for native (PDB ID:5B0H) and ALECT2 fibril (PDB ID: 9NON) were provided as input.

### Figure Panels

All figure panels were created with Adobe Illustrator.

## Acknowledgements

We dedicate this work to the memory of the late Dr. Merrill D. Benson, whose pioneering contributions advanced the understanding of amyloid diseases and brought hope and support to countless affected families. We are deeply grateful to the patients and their families for their generous tissue donations and to Indiana University for providing essential material for this research.

We extend our appreciation to all members of the Structural Biology Laboratory (SBL) and the staff of the UTSW Cryo-Electron Microscopy Facility and Electron Microscopy Core Facility for their technical assistance and access to instrumentation. The SBL and CEMF at UT Southwestern are partially funded by the Cancer Prevention & Research Institute of Texas (CPRIT; grant RP220582). The authors thank the UTSW Proteomics Core for assistance with proteomics analysis and the UTSW Histopathology core for their assistance with histological experiments. We also acknowledge the national cryo-EM center at Stanford-SLAC (project CA172) for providing access to state-of-the-art instrumentation, technical support, and data collection services. All cryo-EM data for this study were acquired at the Stanford-SLAC Cryo-EM Center (S2C2), which is funded by the National Institute of General Medical Sciences (1R24GM154186). The views expressed here are solely those of the authors and do not necessarily reflect the official policies of the National Institutes of Health. We are especially grateful to Lisa B. Dunn and Dr. Alexandre Cassago at S2C2 for their invaluable support and assistance throughout data collection.

Computational analyses were made possible in part by the BioHPC high-performance computing facility at the Lyda Hill Department of Bioinformatics, UT Southwestern Medical Center (https://portal.biohpc.swmed.edu).

Molecular visualization and structural analyses were conducted using UCSF ChimeraX, developed by the Resource for Biocomputing, Visualization, and Informatics at the University of California, San Francisco, with support from NIH R01-GM129325 and the Office of Cyber Infrastructure and Computational Biology, National Institute of Allergy and Infectious Diseases.

We also acknowledge the use of ChatGPT (OpenAI, 2025) to assist with textual refinement and manuscript editing. The authors reviewed and verified all AI-assisted content.

We thank Dr. Michael R. Sawaya, University of California, Los Angeles for sharing his structural expertise.

## Author contributions

Conceptualization: L.S, S.A Experiment design: B.N, V.S, S.A

Data collection: B.N, V.S, S.A, P.B, P.S, M.P, B.E, C.L, Y.A, L.L, R.K, A.L, B.L.B

Data analysis: S.A, B.N, V.S, L.S, C.A Supervision: L.S

Funding acquisition: L.S Manuscript writing: S.A, L.S, V.S Maunscript editing: S.A, L.S, B.N

## Data and Materials Availability

Mass spectrometry data have been deposited to MassIVE database (a member of ProteomeXchange) under accession code MSV000098220. Graphed data is provided in the source data file. Cryo-EM maps have been deposited in the Electron Microscopy Data Bank under accession codes as following; single protofilament morphology (EMD-49601), double protofilament 1 morphology (EMDB-49624), double protofilament 2 morphology (EMDB-49623). The atomic model of ALECT2 single protofilament morphology is available at the Protein Data Bank under accession code 9NON. All data generated or analyzed during this study that support the findings are available within this published article and its supplementary data files. Tissue samples were obtained from the laboratory of the late Dr. Merrill D. Benson at Indiana University. These specimens are under a material transfer agreement with Indiana University and cannot be distributed freely.

