## Supplemental Figures and Table for "Structural polymorphism of *ex-vivo* ALECT2 amyloid fibrils revealed by cryo-EM"

**Affiliations:**
*[1]Center for Alzheimer's and Neurodegenerative Diseases, Department of Biophysics, Peter O'Donnell Jr Brain Institute, University of Texas Southwestern Medical Center (UTSW), Dallas, TX, USA.*
*[2]SciKonnect and BioPatriKa, India.*
*[3]Department of Pathology, University of Texas Southwestern Medical Center (UTSW), Dallas, TX, USA.*
*[4]Department of Ophthalmology, University of Texas Southwestern Medical Center (UTSW), Dallas, TX, USA.*
*[5]Department of Internal Medicine, University of Texas Southwestern Medical Center (UTSW), Dallas, TX, USA.*
*[6]Jerry H. Hodge School of Pharmacy, Texas Tech University Health Sciences Center, Dallas, TX, USA.*
*[7]Department of Biochemistry, University of Texas Southwestern Medical Center, Dallas, TX, USA.*
*[8]The University of New Mexico School of Medicine, Albuquerque, New Mexico, USA.*
*[9]Department of Pathology and Laboratory Medicine, Indiana University School of Medicine, Indianapolis, IN, USA.*

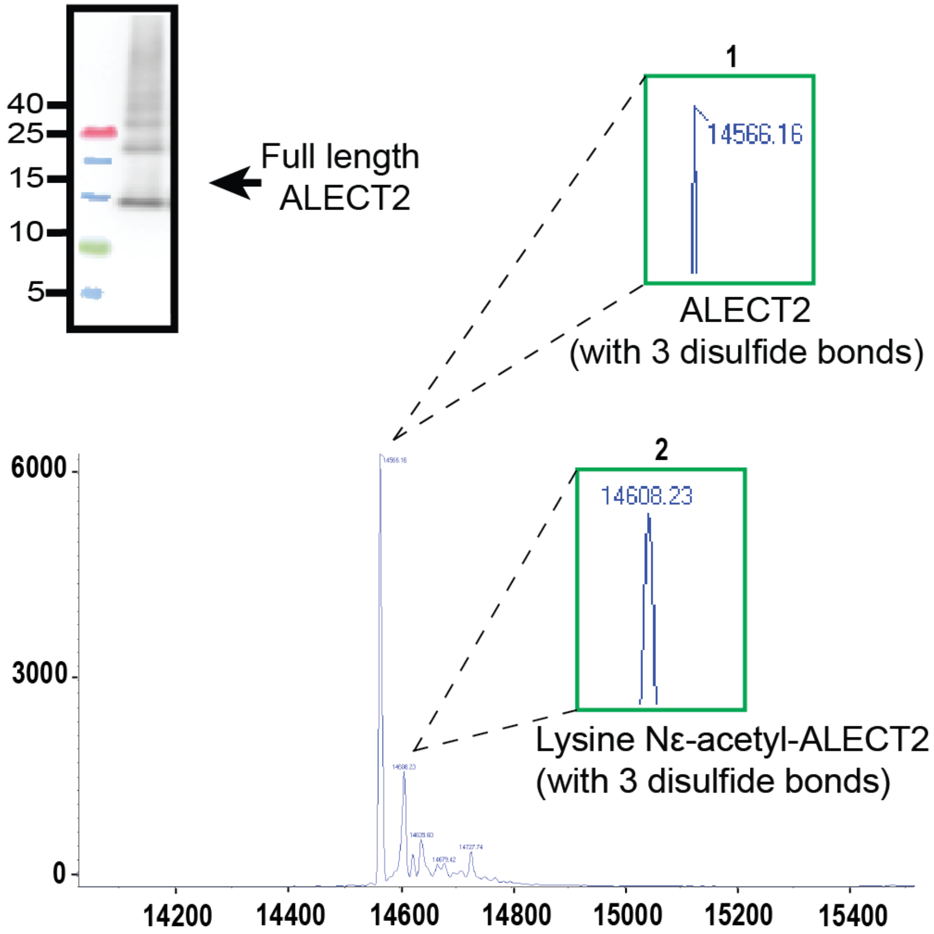

**Supplementary Fig. 1 Characterization of ALECT2 fibrils extracted from the kidney. a** Western blot of the water-soluble fibril elutions from the kidney using an antibody raised against full-length LECT2 protein. Arrow indicates band corresponding to full-length LECT2. Results for **a** representative of n=1 technical replicate. **b.** Intact Mass Analysis of ALECT2 fibrils extracted from the kidney**.** Deconvoluted mass spectrum of ALECT2 fibrils, confirming the presence of full-length ALECT2 with three disulfide bonds (peak 1). We detected additional post-translational modification (acetylation, peak 2) based on mass shifts in the spectrum.

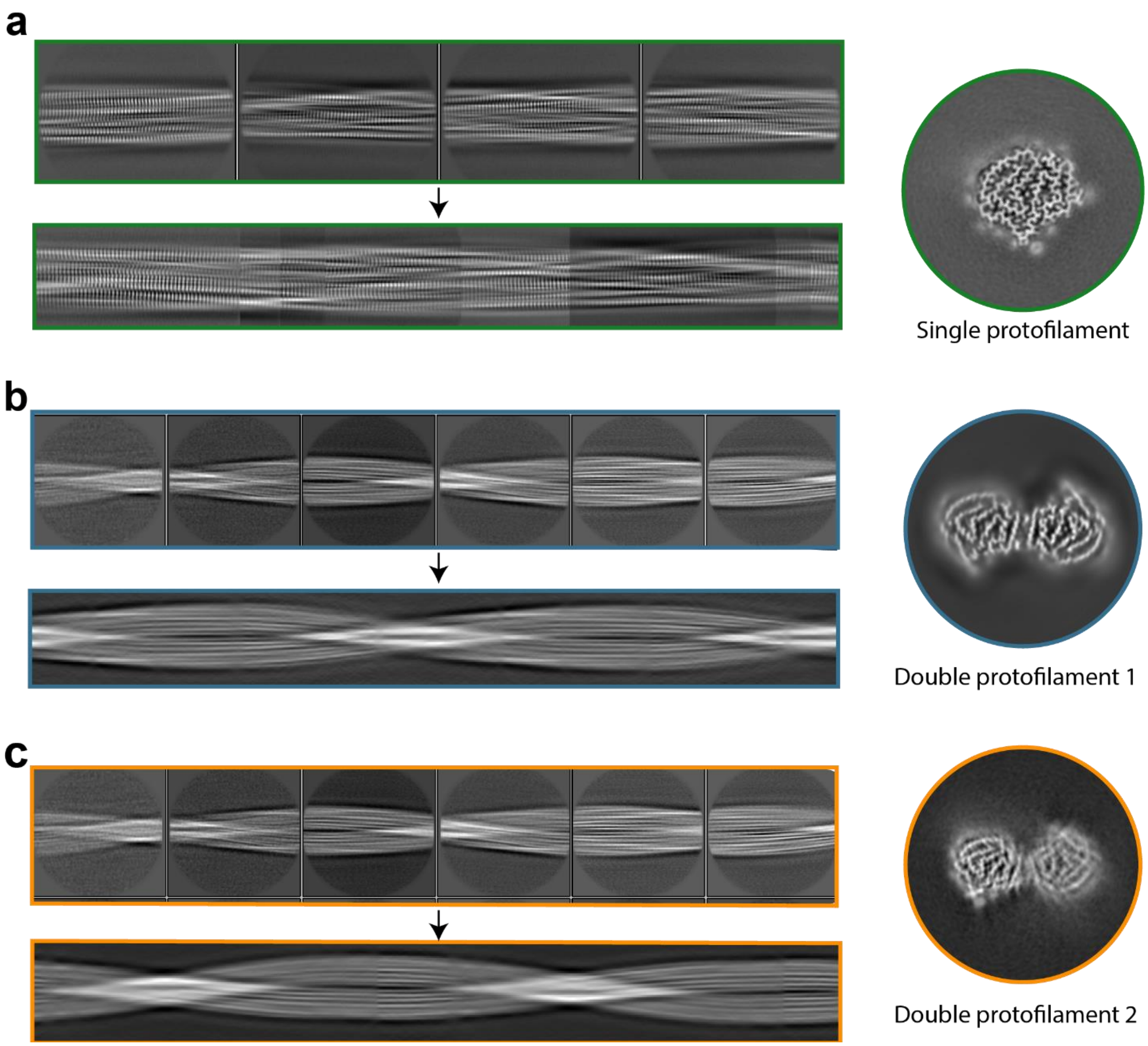

**Supplementary Fig. 2. Representative 2D class averages and 3D reconstructions of ALECT2 single and double protofilament fibrils**. **a.** single protofilament fibrils and **b.** double protofilament 1 fibrils and **c.** double protofilament 2 fibrils, extracted with a particle box size of 352 pixels. Below the individual 2D class panels, we present a stitched view of all 2D classes for the single protofilament fibril and the initial models of the two double protofilament morphologies, generated in RELION.

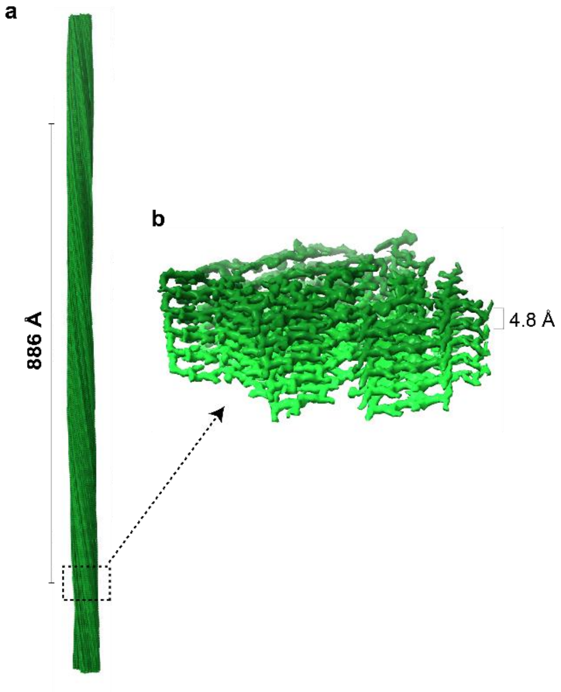

**Supplementary Fig. 3**. **Structural features of the ALECT2 single protofilament highlighting its helical parameters. a.** Side view of the reconstructed ALECT2 single protofilament model showing the crossover distance. **b.** The closeup side view of the map depicting the helical rise and layers.

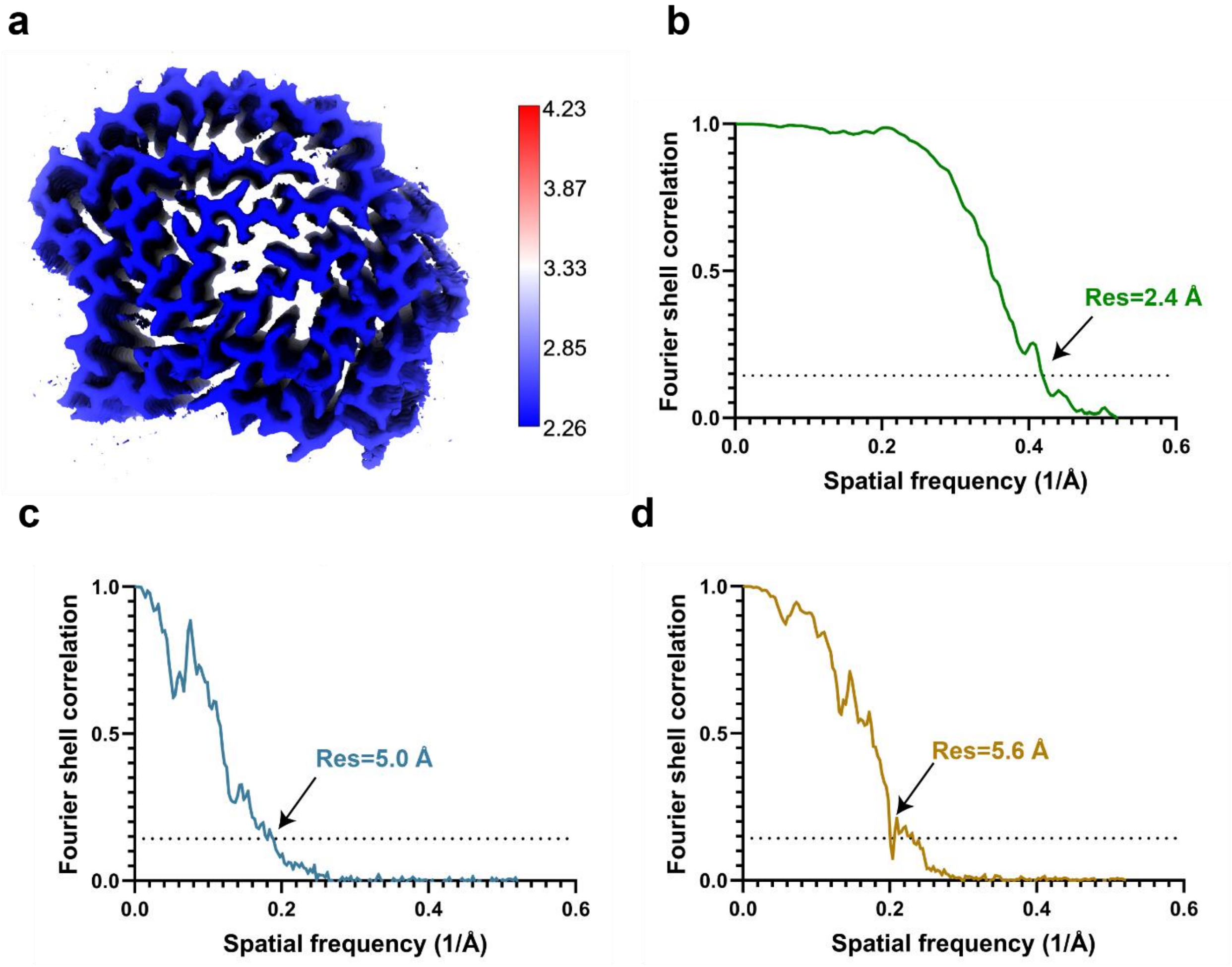

**Supplementary Fig. 4. Resolution Assessment of the Cryo-EM Structure of ALECT2 fibrils from the kidney. a.** Local resolution map of the ALECT2 fibril structure, highlighting variations in resolution across different regions of the density map. Fourier shell correlation (FSC) curve between two independently refined half-maps, illustrating the overall resolution of the reconstruction for ALECT **b.** single protofilament morphology, **c.** Double protofilament 1 morphology and **d.** Double protofilament 2 morphology.

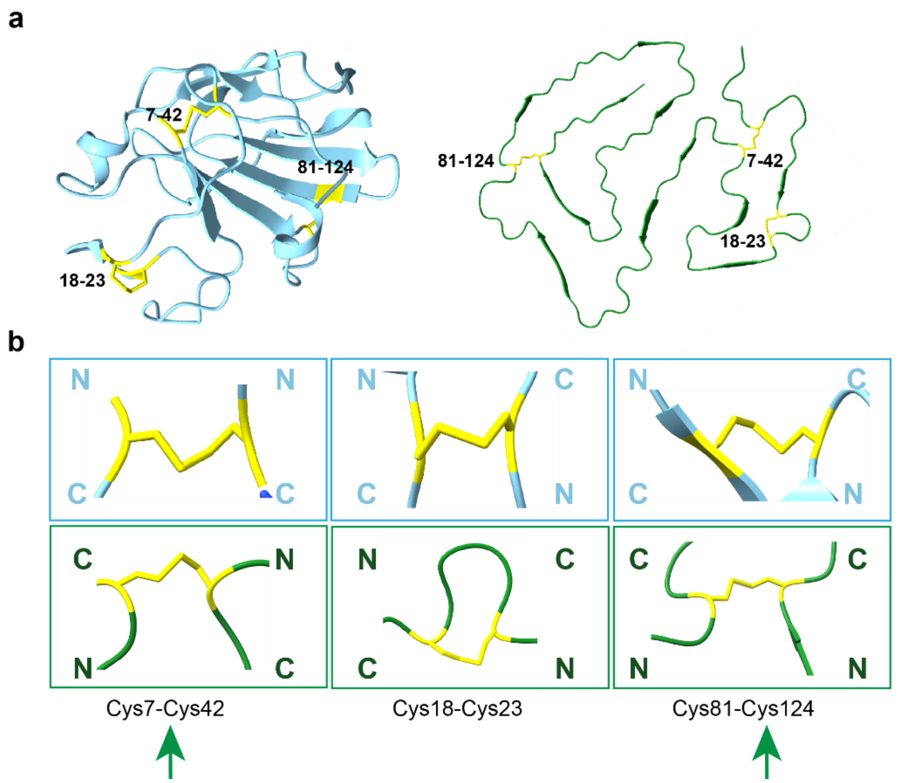

**Supplementary Fig. 5. Structural reorientation of disulfide bonds in ALECT2 fibrils. a.** Secondary structure visualization of native LECT2 monomer from the crystal structure (PDB ID: 5B0H, left) and ALECT2 fibrils from the kidney (right). Disulfide bonds are highlighted in yellow. **b.** Green arrows mark the disulfide bonds that change orientation. Cys7–Cys42 bond, which adopts a parallel N-to-C orientation in the native structure (light blue), shifts to an anti-parallel orientation in the fibril structure. In contrast, the Cys18–Cys23 bond maintains the same orientation in both native and fibrillar states. The Cys81–Cys124 bond transitions from an anti-parallel orientation in the native structure to a parallel orientation in the fibril.

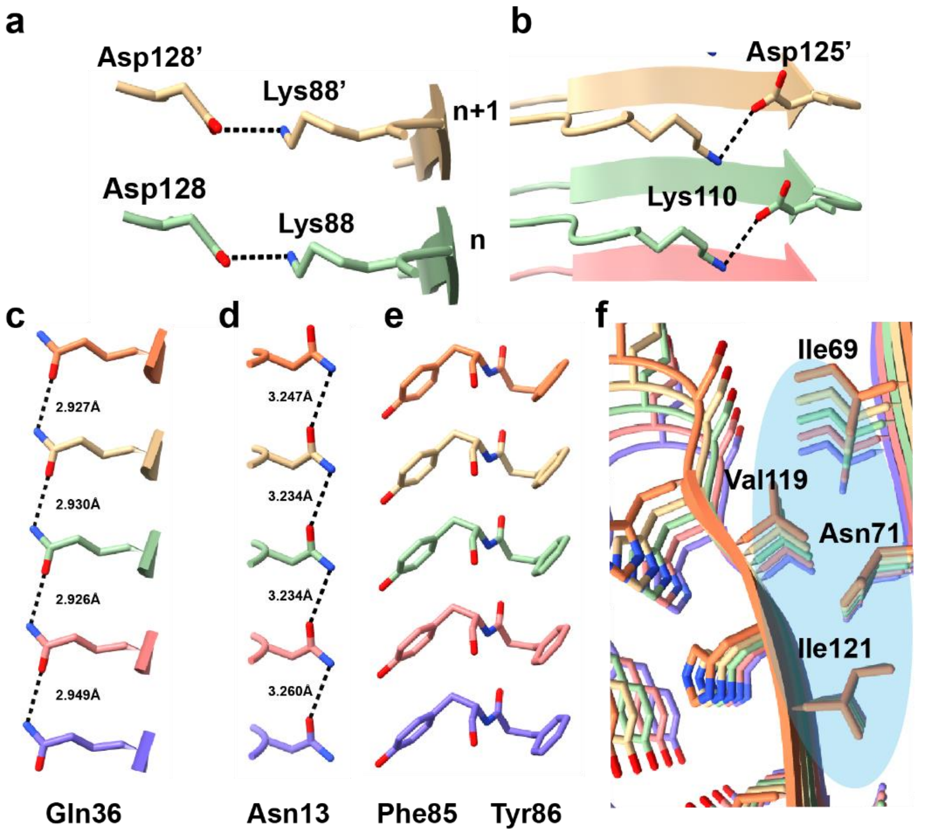

**Supplementary Fig. 6. Additional interactions stabilizing the ALECT2 fibril structure**.

**a and b.** Intra-layer salt bridge between Asp128 and Lys88 **(a),** and between Asp125 and Lys110 **(b)**. **c and d.** Network of hydrogen-binding polar ladders formed by Asn **(c)** and Gln **(d)** residues across all layers **e**. π-π interactions by aromatic residues across all layers represented here by Tyr 86 and Phe 85. **f**. Illustration of steric zippers stabilizing the fibril fold.

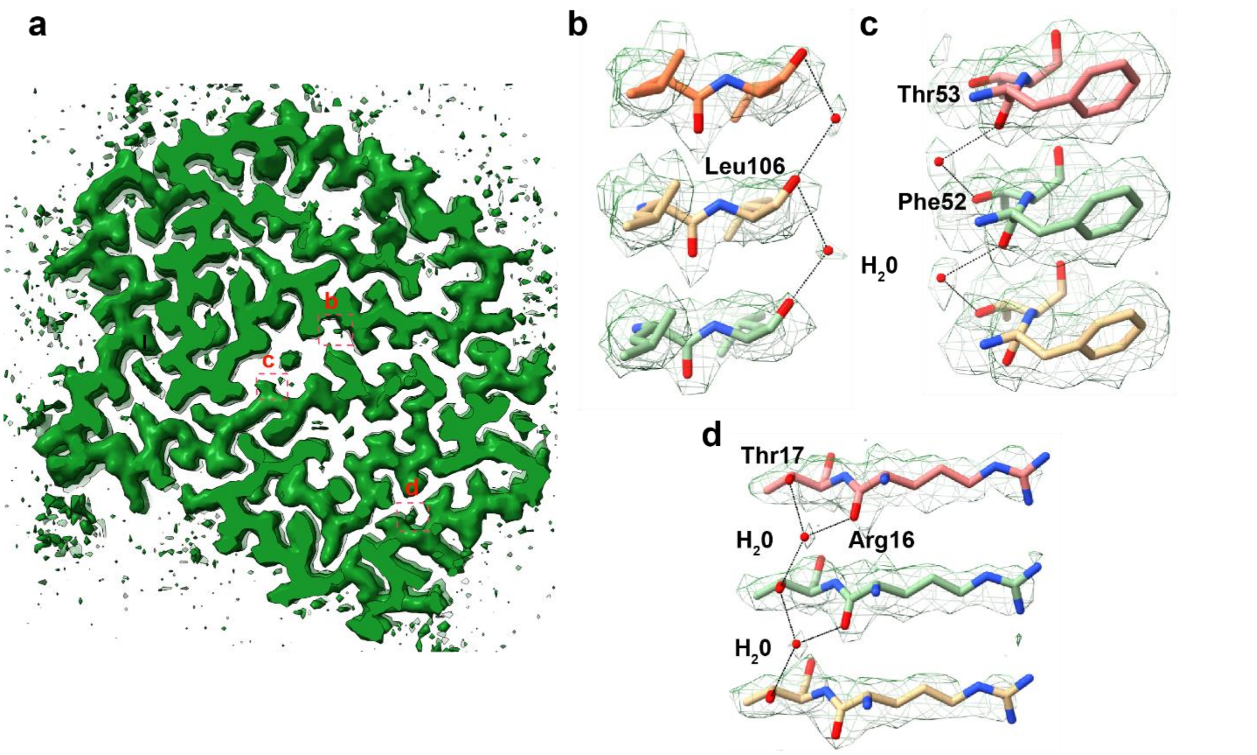

**Supplementary Figure 7**. **Predicted water molecules within the ALECT2 fibril structure**. **a.** ALECT2 map with predicted water molecules marked in red dotted squares. **b.** Water molecules interacting with Leu106 across the fibril layers. **c.** water molecule interacting with Thr53 and Phe52 across the layers and **d.** water molecule interacting with Thr17 and Arg16 across the fibril layers.

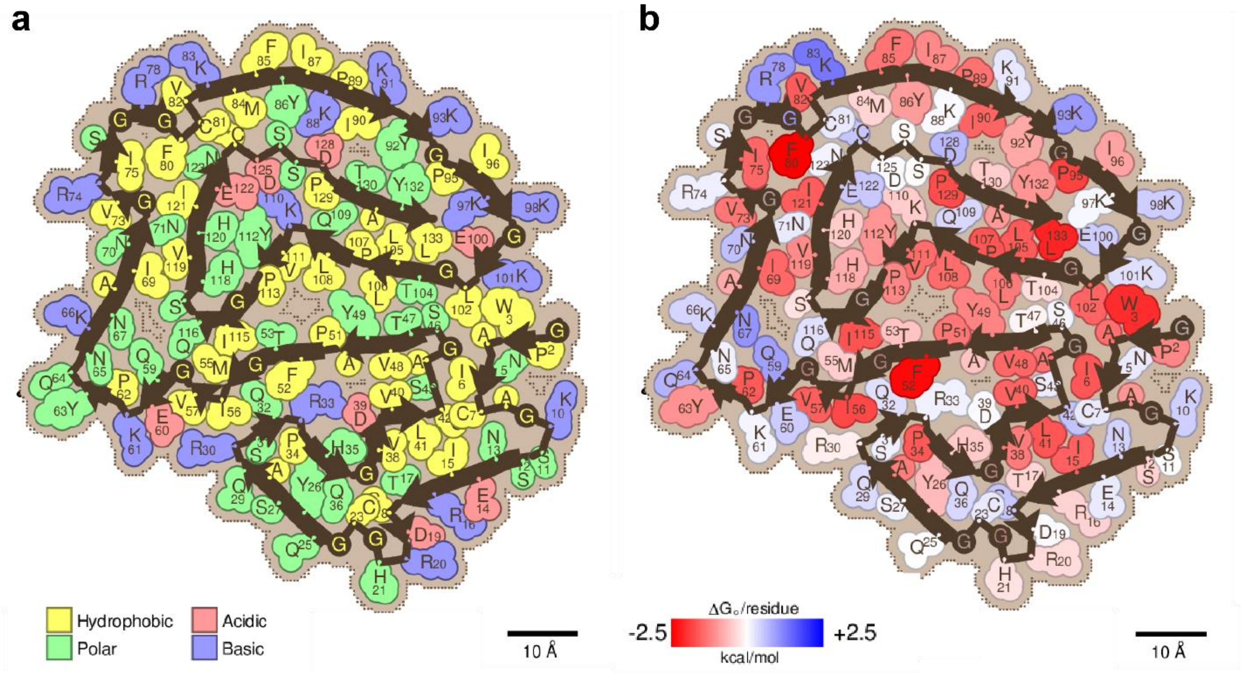

**Supplementary Fig. 8. Composition and stability of the ALECT2 fibril core. a**. Schematic view of ALECT2 fibrils showing residue composition where residues are color coded by amino acid category, as labeled. **b**. Representation of ALECT2 fibril core depicting stabilizing residues. Strongly stabilizing side chains are colored red, and destabilizing side chains are colored blue.

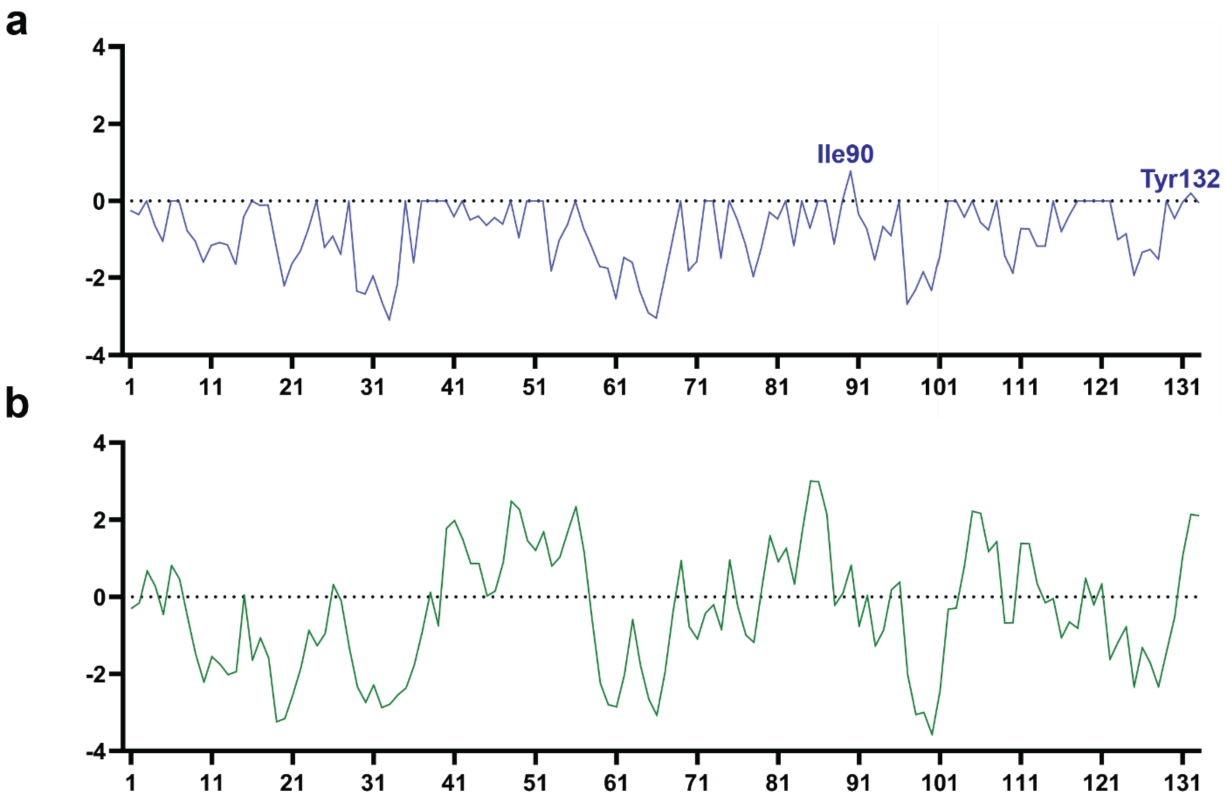

**Supplementary Fig. 9. Comparative analysis of aggregation-prone regions (APR) in LECT2 using AggreScan3D 2.0. a.** AggreScan3D analysis of the native LECT2 monomer (Chain B, PDB: 5B0H). **b.** Aggrescan3D analysis of the ALECT2 fibril structure (Chain C, PDB: 9NON) from this study. The plots display per-residue aggregation scores, where positive values indicate regions prone to aggregation, and negative values suggest aggregation-resistant regions.

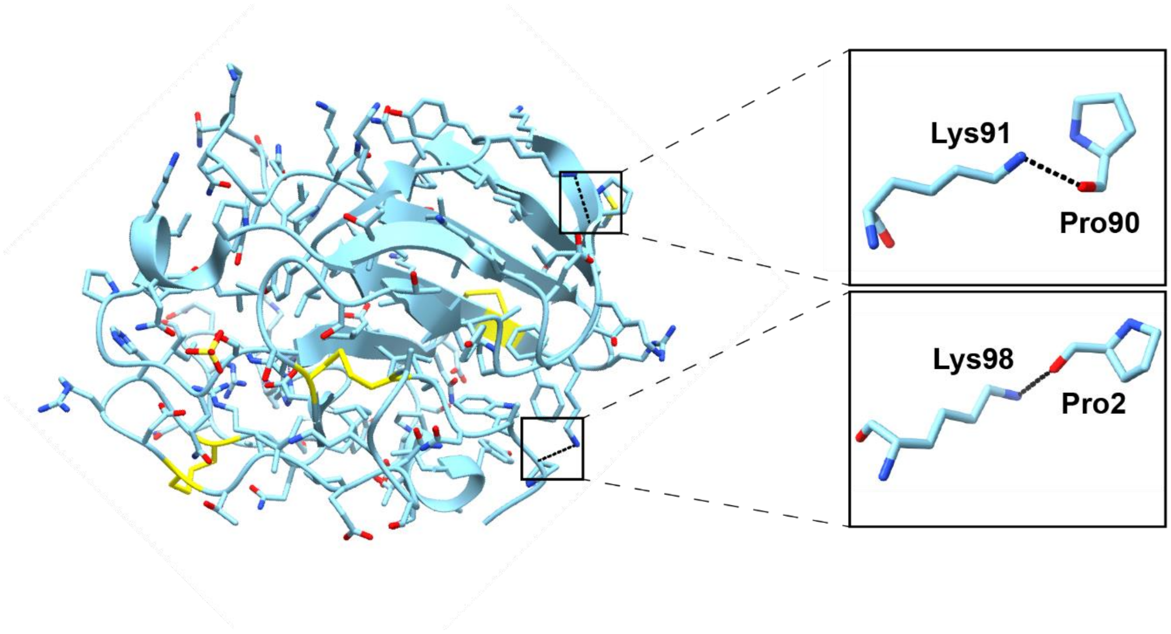

**Supplementary Fig. 10. Interaction between lysine residues and the amide backbone of proline in the native LECT2 structure (Chain B, PDB: 5B0H).** These interactions help stabilize the native fold and would be disrupted by lysine acetylation, potentially destabilizing the structure and promoting aggregation. Dashed lines, distances lower than 3 Å.

a

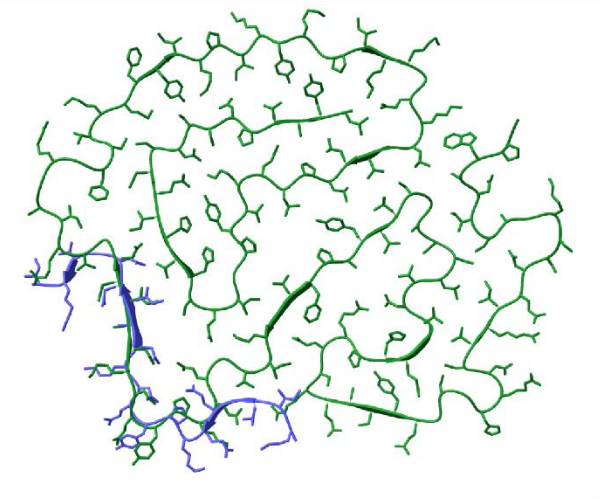

b

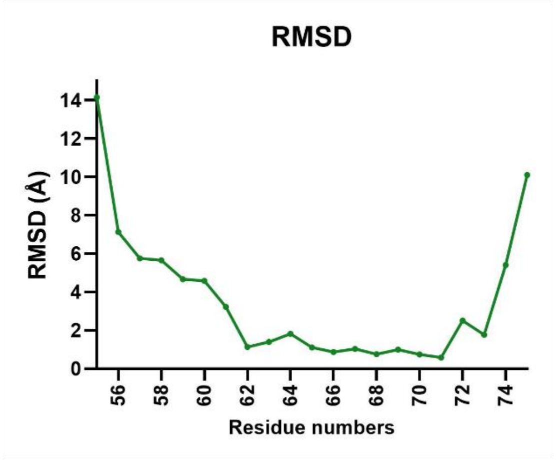

**Supplementary Fig. 11. Structural comparison between recombinant LECT2 and *ex-vivo* single protofilament ALECT2 structure from this study. a.** *Ex-vivo* single protofilament fibril structure (green, PDBID: 9NON) aligned with the corresponding region of the single protofilament of the recombinant LECT2 structure (blue, PDBID: 8G2V). **b.** RMSD comparison between ALECT2 *ex-vivo* and *in-vitro* fibril structure**.**

### Supplementary tables

| Peak | Position | Theoretical $M_w$ (Da) | Observed $M_w$ (Da) | PTMs |
|---|---|---|---|---|
| 1 | 1-133 | 14566.5 | 14566.16 | 3x disulfide bonds |
| 2 | 1-133 | 14608.5 | 14608.23 | 3x disulfide bonds<br>1x acetylation |

**Supplementary Table 1**. Intact Mass spectrometry peaks from Supplementary Figure 1 tabulated. $M_w$, molecular weight. PTMs, post-translational modifications.

| Sequence | Modification | Sequence position in protein | Modified residue |
|---|---|---|---|
| [Y].QNKNAINNGVRISGRGF.[C] | 1xAcetyl | [64-80] | K66 |
| [F].YIKPIKY.[K] | 1xAcetyl | [86-92] | K91 |
| [Y].KGPIKKGEKL.[G] | 1xAcetyl | [93-102] | K98 |
| [F].TGMIVGQEKPY.[Q] | 1xAcetyl | [53-63] | K61 |
| [Y].KGPIKKGEKL.[G] | 1xAcetyl | [93-102] | K101 |

**Supplementary Table 2.** Chymotrypsin-digestion based PTM analysis of ALECT2 fibrils extracted from the kidney reveals multiple lysine acetylation sites.

**Supplementary Table 3: Cryo-EM data collection, refinement and validation statistics**

| | Single (EMD-49601) (PDB 9non) | Double protofilament 1 (EMDB-49624) | Double protofilament 2 (EMDB-49623) |
|---|---|---|---|
| **Data collection and processing** | | | |
| Magnification | | 130,000x | |
| Voltage (kV) | | 300 | |
| Electron exposure (e–/Å$^2$) | | 40 | |
| Exposure Time (sec) | | 3.69 | |
| Number of frames | | 40 | |
| Defocus range (μm) | | -0.8 to -2.6 | |
| Pixel size (Å) | | 0.954 | |
| Symmetry imposed | | C1 | |
| Micrographs collected | | 22,899 | |
| Micrographs processed | 14,112 | 11,943 | 11,943 |
| Total particles (box size 300 px) | 3,802,894 | 3,802,894 | 3,802,894 |
| Particles used in final map | 36,221 | 41,111 | 12,405 |
| Map resolution (Å) | 2.4 | 5.0 | 5.6 |
| FSC threshold | 0.143 | 0.143 | 0.143 |
| Map resolution range (Å) | 2.26 – 2.85 | n/a | n/a |
| **Refinement** | | | |
| Model resolution (Å) | 2.1 | | |
| FSC threshold | 0.143 | | |
| Model resolution range (Å) | 2.1 – 2.4 | | |
| Map sharpening *B* factor (Å$^2$) | 64.76 | | |
| Model composition | | | |
| Chains | 5 | | |
| Non-hydrogen atoms | 0 | | |
| Protein residues | 665 | | |
| Ligands | 0 | | |
| *B* factors (Å$^2$) | | | |
| Protein | 43.29 | | |
| Ligand | 0 | | |
| R.m.s. deviations | | | |
| Bond lengths (Å) | 0.004 | | |
| Bond angles (°) | 1.004 | | |
| Validation | | | |
| MolProbity score | 1.61 | | |
| Clashscore | 2.95 | | |
| Poor rotamers (%) | 0.90 | | |
| Ramachandran plot | | | |
| Favored (%) | 90.84 | | |
| Allowed (%) | 9.16 | | |
| Disallowed (%) | 0.00 | | |
